# Lifespan Neurodegeneration Of The Human Brain In Multiple Sclerosis

**DOI:** 10.1101/2023.03.14.532535

**Authors:** Pierrick Coupé, Vincent Planche, Boris Mansencal, Reda A. Kamroui, Ismail Koubiyr, José V. Manjon, Thomas Tourdias

## Abstract

**Background:** Atrophy related to Multiple Sclerosis (MS) has been found at the early stages of the disease. However, the archetype dynamic trajectories of the neurodegenerative process, even prior to clinical diagnosis, remain unknown.

**Methods:** We modeled the volumetric trajectories of brain structures across the entire lifespan using 40944 subjects (38295 healthy controls and 2649 MS patients). Then, we estimated the chronological progression of MS by assessing the divergence of lifespan trajectories between normal brain charts and MS brain charts.

**Results:** Chronologically, the first affected structure was the thalamus, then the putamen and the pallidum (3 years later), followed by the ventral diencephalon (7 years after thalamus) and finally the brainstem (9 years after thalamus). To a lesser extent, the anterior cingulate gyrus, insular cortex, occipital pole, caudate and hippocampus were impacted. Finally, the precuneus and accumbens nuclei exhibited a limited atrophy pattern.

**Conclusion:** Subcortical atrophy was more pronounced than cortical atrophy. The thalamus was the most impacted structure with a very early divergence in life. It paves the way toward utilization of these lifespan models for future preclinical/prodromal prognosis and monitoring of MS.

## Introduction

Multiple Sclerosis (MS) is a chronic, demyelinating and inflammatory pathology of the central nervous system, that involves significant neurodegenerative damages. The brain atrophy resulting from neurodegeneration is an important biomarker of disease progression, even better than the traditional white matter lesion assessment [1].

Magnetic resonance imaging (MRI) has proven to be a useful tool for estimating brain atrophy in neurodegenerative diseases. In MS, MRI mainly focuses on the white matter lesions to establish the initial diagnosis [2] and to monitor therapeutic response, but MS progression could also be monitored through volumetric measures independently of relapses [3]. Thanks to advances in automatic analysis of brain MRI, the volume of brain structures can be estimated accurately and robustly [4]. Therefore, while previously focusing on global brain volume, recent studies have been able to analyze MS-related brain atrophy more finely at the structural level [5].

Evidence suggests that global brain atrophy is mainly driven by gray matter (GM) alterations rather than to white matter damage [6]. These GM alterations have been found in both cortical and subcortical structures in MS patients [7]. Also, some brain structures seem more likely to be affected than others [5]. GM atrophy has been observed across all the stages of the disease even in preclinical MS stage (*e.g.,* clinical isolated syndromes – CIS) [8]. Moreover, several studies investigated the progression of GM atrophy across multiple sclerosis phenotypes [5]. However, so far, we do not know when such a process starts and the full dynamic over several decades has not been revealed. Without a database starting long before the appearance of the first symptoms and providing the corresponding follow-up over decades, such a study is very challenging.

Recently, the concatenation of a large number of cross-sectional data has been used to reveal the typical course of brain volumes during the lifespan [9]. We pioneered an approach that consists in comparing such normative trajectories with those from patients, and we demonstrated its relevance to estimate preclinical GM atrophy in Alzheimer’s disease [10,11]. To overcome the lack of data prior to clinical diagnosis, pathological lifespan modeling combines healthy subjects and patients covering the entire lifespan. In this study, we propose to adapt this strategy to MS. Thanks to these lifespan models, we present new insights on the spatiotemporal evolution of GM atrophy across the entire lifespan. Moreover, we estimate the most atrophic structures, the sequence of impacted structures and the average age of onset of significant atrophy.

## Methods

### Standard protocol approvals, registrations, and patient consents

All the used images were obtained from public datasets. Database providers ensured compliance with ethical guidelines such as informed consent and anonymization (see Acknowledgments).

### Datasets

In our study, lifespan models of brain volume trajectories were estimated using 24 open-access MRI databases. To this end, we collected MRIs of 41671 subjects, 38978 from cognitively normal, healthy control (HC) subjects covering the entire lifespan (from 1 to 100 years of age) and 2693 from patients with MS. All the MRIs were collected on 1.5T or 3T magnets.

#### HC database

For HC subjects, we used the baseline T1-weigthed MRI from the following datasets: UKbiobank (n=29932, https://www.ukbiobank.ac.uk/), *C-MIND* (n=236, https://research.cchmc.org/c-mind/), *NDAR* (n=382, https://ndar.nih.gov), *ABIDE* (n=492 http://fcon_1000.projects.nitrc.org/indi/abide/), *ICBM:* (n=294 http://www.loni.usc.edu/ICBM/), *IXI* (n=549, http://brain-development.org/ixi-dataset/), *ADNI1&2* (n=404, http://adni.loni.usc.edu), *AIBL* (n=232, http://www.aibl.csiro.au/), OASIS (n=298, https://www.oasis-brains.org), ADHD-200 (n=544, http://fcon_1000.projects.nitrc.org/indi/adhd200/), DLBS (n=315, http://fcon_1000.projects.nitrc.org/indi/retro/dlbs.html), ISYB (n=213, https://www.scidb.cn/en/detail?dataSetId=826407529641672704), MIRIAD (n=23, https://www.ucl.ac.uk/drc/research/research-methods/minimal-interval-resonance-imaging-alzheimers-disease-miriad), PPMI (n=166, https://www.ppmi-info.org/), PREVENT-AD (n=307, https://openpreventad.loris.ca/), Amsterdam open MRI collection (AOMIC_ID100 & PIOP1 & PIOP2, n=1361, https://nilab-uva.github.io/AOMIC.github.io/), Calgary cohort (n=267, https://osf.io/axz5r/), CamCAN (n=653, https://camcan-archive.mrc-cbu.cam.ac.uk/dataaccess/), PIXAR (n=155, https://openneuro.org/datasets/ds002228/versions/1.1.0), SALD (n=494, http://fcon_1000.projects.nitrc.org/indi/retro/sald.html), SRPBS (n=791, https://bicr-resource.atr.jp/srpbsopen/), NACC (n=161, https://naccdata.org), NIFD (n=135, https://memory.ucsf.edu/research-trials/research/allftd) and SLIM (n=574, http://fcon_1000.projects.nitrc.org/indi/retro/southwestuni_qiu_index.html). After quality control (QC – see **Image Processing** section), this database contained 38295 HC subjects (see Table 1).

**Table 1:**
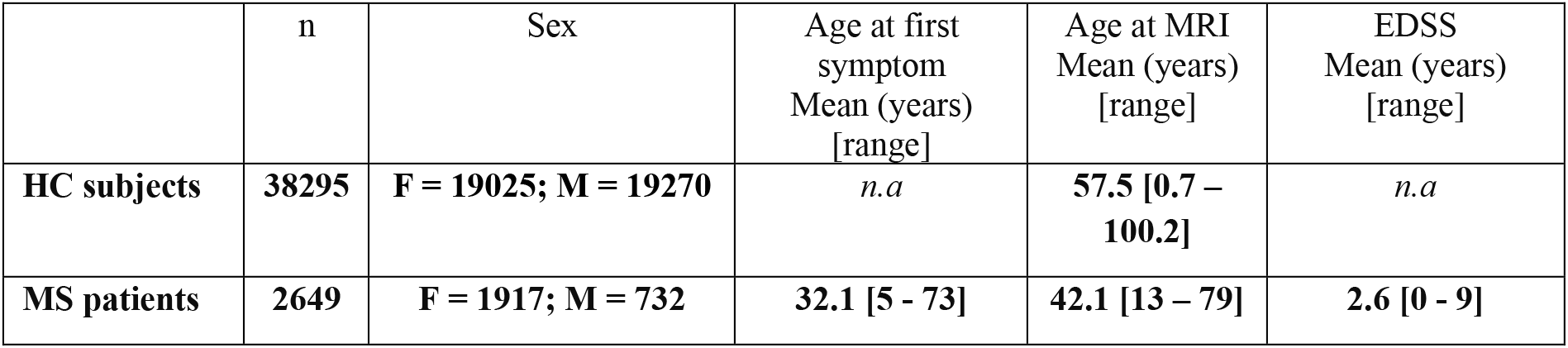
Databases description. This table provides the total number (n) of considered images (after quality control), the gender proportion, and the average ages and intervals in brackets.

#### MS database

For MS patients, we used the first available T1-weighted and FLAIR MRIs from the “Observatoire Français de la Sclérose en Plaques” (OFSEP) dataset (n=2692, https://www.ofsep.org/en/) including 544 subjects with a clinically isolated syndrome (CIS), 1686 patients with relapsing-remitting multiple sclerosis (RRMS), 288 secondary-progressive multiple sclerosis (SPMS) and 174 patients with primary-progressive multiple sclerosis (PPMS) [12]. After QC, this database contained 2649 MS subjects (see Table 1). The average age at the first of symptoms suggestive of MS was 32y and the average age at the MRI used for analysis was 42y.

### Construction of lifespan groups

To study the brain volumetric trajectories of HC subjects and MS patients across the entire lifespan, we compiled several open-access databases to construct normal and diseased models. For the HC models, we used the 38295 MRIs from HC subjects remaining after QC covering the entire lifespan (see Table 1). For the MS models, we followed the strategy proposed in [10,11] for lifespan analysis of Alzheimer’s disease. This framework is based on the assumption that neurodegeneration is a continuous and progressive process along the pathology evaluation. Therefore, to constrain the model over the entire lifespan, it has been proposed to mix HC with patients. Herein, we combined MRIs of 2649 MS patients after QC (see Table 1) with MRIs of 3711 healthy controls younger than 23 years. This age was the quantile at 5% of the MS population that enabled a smooth transition from HC subjects to MS patients. These HC subjects were all the subjects younger than 23y in the 38295 HC subjects used for HC models after QC. Consequently, for the MS models, between 1y-13y only HC subjects were used, between 13y-23y a mix of HC subjects and MS patients were used, and after 23y only MS patients were used. At the end, the parametric MS models were constrained over the entire lifespan using 6360 subjects. To ensure model validity, we analyzed results over 1-62y, 62y being the quantile at 95% of the MS population age distribution.

### Image processing

All the considered T1-weighted MRI were segmented using AssemblyNet [13] (https://github.com/volBrain/AssemblyNet/). This software produce sfine-grained segmentation (*i.e.,* 132 structures) of the entire brain.

#### Pipeline for HC subjects

All the T1-weighted MRI were first preprocessed to harmonize them. The preprocessing consisted of denoising [14], inhomogeneity correction [15], affine registration into the Montreal Neurological Institute (MNI) space [16], tissue-based intensity normalization [17] and intracranial cavity segmentation [18]. Afterward, all the preprocessed images were checked by automatic quality control (QC) based on artificial intelligence [19]. Finally, structure segmentation was achieved using 250 deep learning models through a multiscale framework [13].

#### Pipeline for MS patients

For MS patients, T1-weighted MRI were preprocessed as for HC. In addition, the FLAIR images were processed using denoising [14], inhomogeneity correction [15], rigid registration into the T1-weighted MRI native space and then projection into Montreal Neurological Institute (MNI) space using T1-weighted registration matrix. Afterward, MS lesions were segmented using DeepLesionBrain [20] (https://github.com/volBrain/DeeplesionBrain/). The lesion masks were then used to perform in-painting of MS lesions on T1-weighted MRI [21]. This step was done to limit the impact of MS lesions visible in T1-weighted MRI on brain segmentation. As for HC, at the end, the preprocessed images were controlled using automatic QC [19] and segmented using AssemblyNet [13].

In the following, we considered 124 structures of the 132 structures produced by AssemblyNet according to the Neuromorphometrics protocol. First, we used 120 symmetric structures (60 left and 60 right): 9 subcortical structures, 17 frontal gyri/lobules, 8 temporal gyri/lobules, 6 parietal gyri/lobules, 8 occipital gyri/lobules, 6 gyri in the limbic cortex, 5 sub-regions of the insular cortex and the cerebellar GM. Moreover, we used four central structures: the brainstem and three groups of vermal cerebellum lobules (*i.e.,* lobules I-V, lobules VI-VII and lobules VIII-X).

### Lifespan trajectory estimation

As in [11], we used normalized volumes (% of total intracranial volume) to compensate for the head size. Afterward, we used a z-score normalization to enable comparison between structures of different sizes. For each structure, the mean and the standard deviation of the volumes over the HC were used to normalize all the volumes (*i.e.,* both normal and pathological).

Moreover, as in [10], we considered different model types to estimate the best trajectory of each brain structure. To this end, we first estimated several low-order polynomial models (*i.e.,* linear model, quadratic model, and cubic model). Then, we kept a model as a potential candidate when F-statistic based on ANOVA (*i.e.*, model vs. constant model) was significant (p<0.05) and all its coefficients were significant using t-statistic (p<0.05). We finally used the Bayesian Information Criterion (BIC) to select the best candidate. This procedure was done for all the structures and for both populations. All the performed statistics were done using Matlab with default parameters.

### Divergence between pathological and healthy models

Once the models were estimated for both populations, distances between healthy and MS trajectories were computed for each brain structure. We used an adjusted confidence level of 95% (*i.e.,* 99.96% after correction for multiple comparisons using Dunn’s procedure) to estimate the prediction bounds of the trajectories. Moreover, we considered that a structure significantly diverged from normal aging when the adjusted 95% confidence intervals of CN models and MS models do not overlap (Fig. 1). Then, all divergent structures were mapped across time and space on the MNI template (see Fig. 2). Finally, the sequence of significant divergence of the most affected brain structures (top 25 structures diverging the most) was listed in chronological order to obtain the MRI staging scheme of lifespan MS atrophy (see Fig. 3).

**Figure 1.**
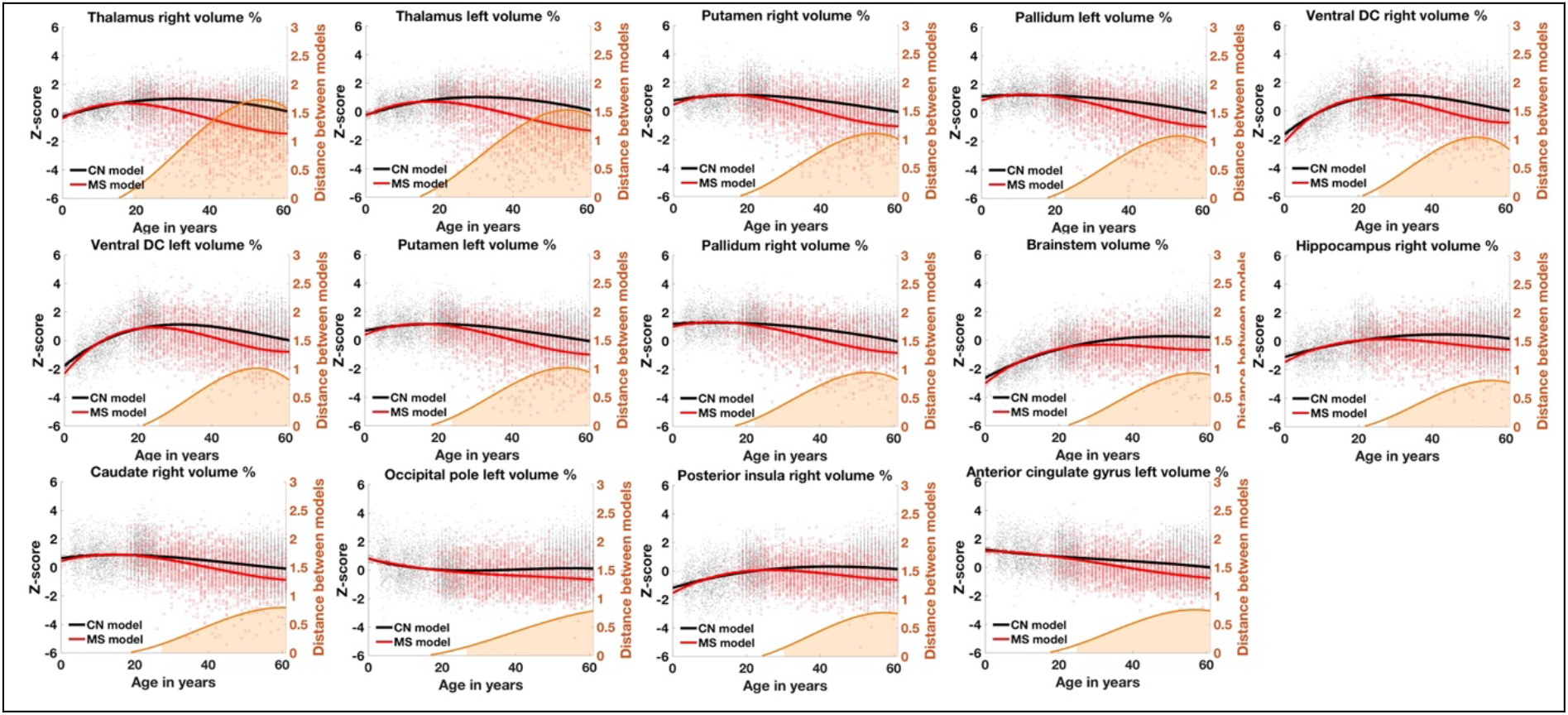
Lifespan trajectories based on z-scores of the main impacted brain structures for healthy aging subjects (in black) and MS patients (in red). Black dots represent all healthy individuals and red dots MS patients. The orange curves represent the distance between the healthy and pathological models. The orange areas indicate the time period where confidence intervals of both models do not overlap. The prediction bounds of the models are estimated with an adjusted confidence level at 95%.

**Figure 2.**
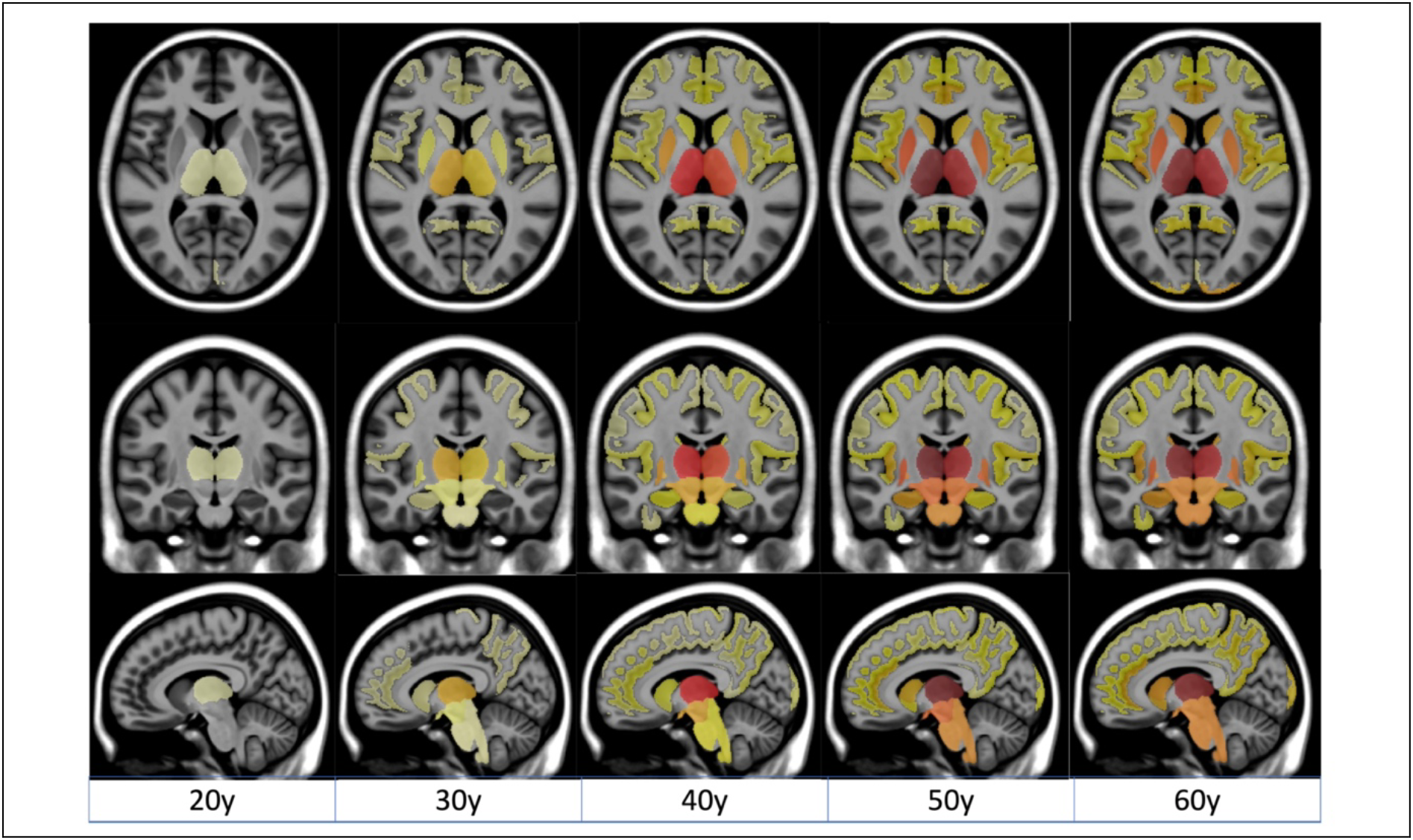
Spatiotemporal progression of the MS-related atrophy. Progression of MS-related atrophy along the three axes with radiological convention for all the structures.

**Figure 3.**
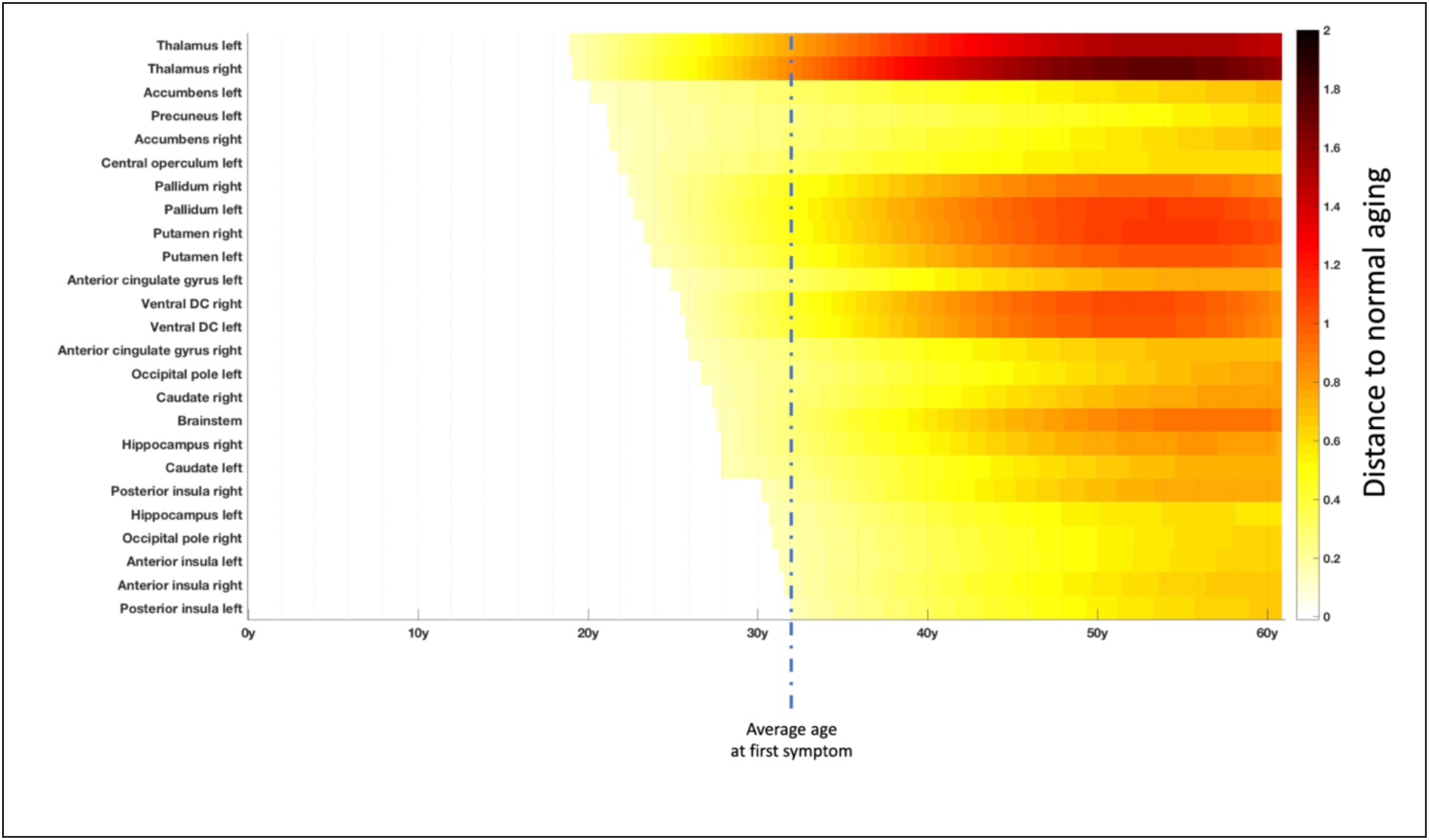
Chronological progression of MS over the most impacted structures. Timeline representing the sequential divergence of the most atrophied structures (top 25) between healthy and MS trajectories. The effectsize of structural divergence is color-coded according to the bar at the bottom right of the figure. The mean age at the first symptom (32y) in this cohort is illustrated with vertical dashed line.

## Results

We found that 33 left brain structures, 32 right brain structures and 3 central structures (*i.e.,* brainstem, and two groups of cerebellar lobules) significantly diverged between MS patients and normal brain charts. Therefore, on the 124 studied structures, we found that 68 of them (more than 50%) exhibited significant smaller volumes for MS trajectories.

### Most impacted structures in terms of atrophy

First, we analyzed the most affected structures over time in terms of atrophy peaks (*i.e.,* maximum distance to the HC models). Figure 1 shows models for the most impacted structures. These structures were the right thalamus (1.72 at 53y), the left thalamus (1.53 at 54y), the right putamen (1.11 at 54y), the left pallidum (1.09 at 53y), the right ventral diencephalon (DC – 1.04 at 52y), the left ventral DC (1.01 at 52y), the left putamen (1.01 at 54y), the right pallidum (0.94 at 53y), the brainstem (0.92 at 57y), the right hippocampus (0.81 at 56y), the right caudate (0.79 at 60y), the left occipital pole (0.78 at 61y), the right posterior insula (0.77 at 57y) and left anterior cingulate (0.76 at 57y).

### Spatiotemporal evolution of atrophy related to MS

Afterward, we studied the spatiotemporal evolution of the MS patients compared to normal brain charts. To this end, we performed a mapping of the divergence between HC and MS trajectories over an MRI atlas (see Fig. 2) and we estimated a global timeline of trajectory divergence considering the top 25 most atrophic structures (see Fig. 3).

Thanks to these analyses, we observed that the most pronounced atrophy related to MS started in deep GM structures (mainly thalamus, pallidum and putamen) then spread through the ventral diencephalon (a structure regrouping the hypothalamus, mammillary body, subthalamic nuclei, substantia nigra, red nucleus, lateral geniculate nucleus, and medial geniculate nucleus) to finally reach the brainstem.

The mean age of the most significant atrophy onset was 19y for the thalamus, between 24-24y for the pallidum and the putamen, around 26y for the ventral diencephalon and 28y for the brainstem. This is to contrast with the mean age of the first symptom that was of 32 years old in our MS cohort. Overall, this means that such lifespan model could help us to estimate atrophy related to MS more than one decade before the first symptom.

Besides, although less important than deep GM atrophy, we also found cortical atrophy started in the parietal lobe (mainly in the precuneus around 21y) and the insular cortex (mainly in the central operculum around 22y) before reaching the limbic cortex (mainly the anterior cingulate gyrus around 25y) to finally end in the occipital lobe (mainly the occipital pole around 27y) and the temporal lobe (mainly the hippocampus around 28y).

## Discussion

In this study, we used a massive number of subjects (N=40944) to model the archetype progression of MS-related atrophy at the structure level across the entire lifespan. Thanks to this modeling, we inferred the spatiotemporal sequence of GM atrophy in MS over the entire course of the disease, including the preclinical stage. Moreover, such framework accounted for atrophy due to normal aging since healthy and diseased lifespan models were compared. Thereby, we automatically estimated the most impacted structures, the atrophy evolution and the average age of atrophy onset.

The MRI staging of atrophy involved the thalamus, then the pallidum and putamen, followed by the ventral diencephalon and finally the brainstem. Moreover, we found that the anterior cingulate gyrus, the insular cortex, and the hippocampus were impacted but on a smaller extent. Finally, we observed that the precuneus and the accumbens nuclei, while early impacted, were slightly atrophic compared to the thalamus. Most of these structures have been previously reported as atrophic in the literature [5,22]. The chronological sequence of atrophy reported here is also consistent with a previous study using an event-based analysis [5]. Our findings also provide new knowledge that could not have been addressed with these previous studies.

Importantly, we estimated the very early divergence in life of normal and pathological brain charts, more than one decade before the average age of symptoms onset (*i.e.,* 32-years-old). This observation echoes the volume loss that was already reported in small sample size cohorts of patient with radiologically and clinically isolated syndrome [8], but also provides an estimation of the mean age of divergence which has never been reported before. While it is well-known that several white matter lesions are already visible at the time of diagnosis (which is the basis for the MRI-based criteria of dissemination in space and time after a first clinical episode [2]), we also demonstrate here that several GM structures will exhibit atrophy at this time. This argues for a neurodegenerative component that is very early and probably compensated for before that lesions would reach eloquent areas leading to a first clinical episode and, in turn, to the clinical diagnosis [23].

The thalamus is the earliest affected structure which is in line with previous literature pointing out the thalamus as a sensitive MRI biomarker of neurodegeneration in the early stage of MS [5,22,24,25]. The thalamus is correlated with a wide range of clinical manifestations and is an important biomarker of disease progression [26]. Significant thalamic atrophy has been found in the early stage of MS suggesting that neurodegeneration begins long before the first symptoms [27]. However, so far, the dynamic of thalamic atrophy at the preclinical stage of MS was unknown. It is likely that thalamic atrophy and atrophy of the other deep gray matter nuclei could be altered through several mechanisms explaining their particular vulnerability early in life of the MS patients [28]. Indeed, it is known that these structures can be altered indirectly through disconnection of their projections by white matter lesions [29]. Direct targeting might add to this secondary phenomenon and therefore accelerate the overall damages. In this process, the high amount of iron within the deep nuclei [30] could accelerate oxydative stress [31]. The deep nuclei adjacent to the CSF of the ventricles could also be directly targeted by inflammatory and neurotoxic soluble factors coming from CSF [32].

On top of the thalamus and the other deep grey matter nuclei, we also found rapid volume loss affecting the brainstem that is directly connected with thalamus and could therefore share the same vulnerability. This also reminds atrophy of the cervical spinal cord that is known to take place rapidly [33]. The cortical ribbon is altered at later stages with some regions showing earlier and more pronounced volume loss than others, especially the hippocampus and the insular cortex. The micro or macrostructural vulnerability of these regions have been continuously highlighted in cross-sectional and short-term longitudinal analyses [5]. Their vulnerability could be related to a large number of connections and therefore a higher probability to be affected by white matter disconnections. The CSF flow is also supposed to be more restricted within the deep invaginations facing these sites which could drive more alterations induced by meningeal inflammation [34].

In order to build such average models, we had to make few assumptions. The first is that there is a smooth transition between the normal and pathological states which is the rational for combining volumes of HC early in life with those from MS patients. This is likely the case regarding (i) the slow rate of atrophy in MS reported before and (ii) the previous validation of our modeling strategy using longitudinal data in other conditions such as Alzheimer’s disease [10,11]. The other assumption is that all the clinical phenotypes of MS follow the same dynamic which is the rational for combing all of them within a single mean model. There are some histological data showing that the pathological processes are regionally consistent between early relapsing-remitting and progressive MS [35]. Furthermore, through other approaches, other authors have reported consistent sequence of events between different clinical phenotypes [5,27].

Overall, the proposed lifespan models could have future potential interests in prognosis and diseased monitoring. We recently showed for Alzheimer’s disease that the distance to healthy and diseased lifespan models can be used to detect neurodegenerative pathologies at their earliest stage while taking into account normal aging [36]. Therefore, as perspectives, such MS brain charts from high number of data modeling the archetype trajectories are likely to be used for comparing individual patient against the mean dynamic profile, at diagnosis or under therapies.

## Acknowledgements

The C-MIND data used in the preparation of this article were obtained from the C-MIND Data Repository created by the C-MIND study of Normal Brain Development. A listing of the participating sites and a complete listing of the study investigators can be found at https://research.cchmc.org/c-mind.

The NDAR data used in the preparation of this manuscript were obtained from the NIH-supported National Database for Autism Research (NDAR). This is supported by the National Institute of Child Health and Human Development, the National Institute on Drug Abuse, the National Institute of Mental Health, and the National Institute of Neurological Disorders and Stroke. A listing of the participating sites and a complete listing of the study investigators can be found at http://pediatricmri.nih.gov/nihpd/info/participating_centers.html.

The ICBM data used in the preparation of this manuscript were supported by Human Brain Project grant PO1MHO52176-11 and Canadian Institutes of Health Research grant MOP-34996.

The IXI data used in the preparation of this manuscript were supported by the U.K. Engineering and Physical Sciences Research Council (EPSRC) GR/S21533/02 - http://www.brain-development.org/.

The ABIDE data used in the preparation of this manuscript were supported by ABIDE funding resources listed at http://fcon_1000.projects.nitrc.org/indi/abide/.

The ADNI data used in the preparation of this manuscript were obtained from the Alzheimer’s Disease Neuroimaging Initiative (ADNI) (National Institutes of Health Grant U01 AG024904). The ADNI is funded by the National Institute on Aging and the National Institute of Biomedical Imaging and Bioengineering and through generous contributions from private partners as well as nonprofit partners listed at: https://ida.loni.usc.edu/collaboration/access/appLicense.jsp. Private sector contributions to the ADNI are facilitated by the Foundation for the National Institutes of Health (www.fnih.org). The grantee organization is the Northern California Institute for Research and Education, and the study was coordinated by the Alzheimer’s Disease Cooperative Study at the University of California, San Diego. ADNI data are disseminated by the Laboratory for NeuroImaging at the University of California, Los Angeles. This research was also supported by NIH grants P30AG010129, K01 AG030514 and the Dana Foundation.

The AIBL data used in the preparation of this manuscript were obtained from the AIBL study of ageing funded by the Common-wealth Scientific Industrial Research Organization (CSIRO; a publicly funded government research organization), Science Industry Endowment Fund, National Health and Medical Research Council of Australia (project grant 1011689), Alzheimer’s Association, Alzheimer’s Drug Discovery Foundation, and an anonymous foundation. See www.aibl.csiro.au for further details.

The ADHAD, DLBS and SALD data used in the preparation of this article were obtained from http://fcon_1000.projects.nitrc.org (Mennes M et al., NeuroImage, 2013; Wei D et al., bioRxiv 2017). The ISYB data were download from https://www.scidb.cn (Imaging Chinese Young Brains, https://doi.org/10.11922/sciencedb.00740).

Data used in the preparation of this article were also obtained from the MIRIAD database (Malone IB et al., NeuroImage, 2012) The MIRIAD investigators did not participate in analysis or writing of this report. The MIRIAD dataset is made available through the support of the UK Alzheimer's Society (Grant RF116). The original data collection was funded through an unrestricted educational grant from GlaxoSmithKline (Grant 6GKC).

Data used in the preparation of this article were obtained from the Parkinson’s Progression Markers Initiative (PPMI) database (www.ppmi-info.org). PPMI – a public-private partnership – was funded by The Michael J. Fox Foundation for Parkinson’s Research and funding partners that can be found at https://www.ppmi-info.org/about-ppmi/who-we-are/study-sponsors.

The Amsterdam open MRI collection AOMIC ID-1000/PIOP1/PIOP2 data used in the preparation of this article were obtained from https://nilab-uva.github.io/AOMIC.github.io/ (Snoek L et al., Scientific data, 2021).

The Calgary preschool MRI dataset was available at https://osf.io/axz5r/ and supported by University of Calgary and CIHR (IHD-134090 & MOP-136797).

Data collection and sharing for this project was provided by the Cambridge Centre for Ageing and Neuroscience (CamCAN, https://camcan-archive.mrc-cbu.cam.ac.uk/dataaccess/). CamCAN funding was provided by the UK Biotechnology and Biological Sciences Research Council (grant number BB/H008217/1), together with support from the UK Medical Research Council and University of Cambridge, UK.

The Pixar database and related fundings were available at https://openneuro.org/datasets/ds000228/versions/1.1.0 (Richardson H et al., Nat Commun, 2018).

Data used in the preparation of this work were obtained from the DecNef Project Brain Data Repository (https://bicr-resource.atr.jp/srpbsopen/) gathered by a consortium as part of the Japanese Strategic Research Program for the Promotion of Brain Science (SRPBS) supported by the Japanese Advanced Research and Development Programs for Medical Innovation (AMED, Tanaka SC et al., Scientific data, 2021).

FTLDNI was funded through the National Institute of Aging, and started in 2010. The primary goals of FTLDNI were to identify neuroimaging modalities and methods of analysis for tracking frontotemporal lobar degeneration (FTLD) and to assess the value of imaging versus other biomarkers in diagnostic roles. The Principal Investigator of NIFD was Dr. Howard Rosen, MD at the University of California, San Francisco. The data are the result of collaborative efforts at three sites in North America. For up-to-date information on participation and protocol, please visit http://memory.ucsf.edu/research/studies/nifd. Data collection and sharing for this project was funded by the Frontotemporal Lobar Degeneration Neuroimaging Initiative (National Institutes of Health Grant R01 AG032306). The study is coordinated through the University of California, San Francisco, Memory and Aging Center. FTLDNI data are disseminated by the Laboratory for Neuro Imaging at the University of Southern California. The NACC database was funded by NIA/NIH Grants listed at https://naccdata.org/publish-project/authors-checklist#acknowledgment.

This research has been conducted using the UK Biobank Resource under application number 80509. See https://www.ukbiobank.ac.uk/ for further details.

Data collection has been supported by a grant provided by the French State and handled by the ”Agence Nationale de la Recherche,” within the framework of the ”Investments for the Future” programme, under the reference ANR-10-COHO-002, Observatoire Français de la Sclérose en Plaques (OFSEP)” & ”Eugène Devic EDMUS Foundation against multiple sclerosis”.

We wish to thank all investigators of these projects who collected these datasets and made them freely accessible. This manuscript reflects the views of the authors and may not reflect the opinions or views of the database providers.

